# Impaired Peri-infarct Long-Term Potentiation Suggests Alternative Mechanisms of Post-stroke Recovery in Rats

**DOI:** 10.1101/2025.10.24.684483

**Authors:** Clément Vitrac, Meret Branscheidt, Wala J Mahmoud, Andreas R. Luft

## Abstract

**Background:** Early after stroke, a period of heightened plasticity in the peri-infarct cortex is thought to provide the physiological substrate for functional motor recovery through increased expression of long-term potentiation (LTP). Prior slice electrophysiology studies on the capacity for LTP after stroke reported conflicting results. Slice preparation could have influenced the results by disrupting neuromodulatory processes altered after stroke. Therefore, whether LTP can be induced in the peri-infarct cortex remains to be elucidated. We investigated LTP in the peri-infarct cortex of anesthetized rats using a novel *in vivo* method that preserves local and long-range circuit dynamics.

**Methods:** LTP, synaptic transmission and short-term plasticity were assessed under urethane anaesthesia in 15 rats (11 with focal stroke to the primary motor cortex, and 4 sham operated controls). Rats were tested one or two weeks after stroke using a minimally perturbed *in vivo* LTP induction protocol validated in naïve rats. Motor function was evaluated using the cylinder test at baseline and at one or two-weeks post stroke.

**Results:** LTP in the peri-infarct cortex was supressed in the stroke group compared to sham controls. In addition, synaptic transmission was reduced for higher stimulation intensities, and short-term plasticity shifted from facilitation to depression, indicating impaired synaptic function at one- and two-weeks post-stroke. Behaviorally, the lesioned rats exhibited motor deficits at one week but showed full recovery by two weeks post-stroke.

**Conclusion:** In recovering animals, both LTP and synaptic transmission are profoundly impaired in the peri-infarct cortex. These findings suggest that mechanisms other than LTP-based plasticity underlie motor recovery during this stage.

## Introduction

The early post-acute stroke period is characterised by spontaneous behavioral recovery that occurs within a narrow time window, during which animals and humansalso exhibit heightened responsiveness to training^1,2^. This time-limited enhanced capacity for behavioral improvement has been attributed to a transient increase in cortical plasticity within the peri-infarct region, particularly through elevated expression of long-term potentiation (LTP). LTP, a process causally linked to learning and memory^3,4^, is supposed to provide a physiological bridge between structural and functional synaptic changes by stabilizing newly formed synapses and enhancing synaptic transmission^3,5^.

The parallels between changes in the peri-infarct cortex early after stroke and those observed during motor learning have lent support to this hypothesis. These include dependence of motor recovery on AMPA and NMDA receptors activation and the production of brain-derived neurotrophic factor^6^, all of which facilitate LTP induction^7,8^. Pioneer work by Hagemann et al. reported increased LTP induction in the sensorimotor peri-infarct cortex using ex-vivo electrophysiology^9^. Following ex-vivo work in the hippocampus has provided evidence of the opposite: a reduced capacity for LTP induction^10,11^. Thus, whether LTP is enhanced or suppressed after stroke remains unresolved.

Importantly, LTP induction in the motor cortex is strongly regulated by local and long-range influences, including the requirement for GABAergic disinhibition^12,13^ and intact dopaminergic transmission from the ventral tegmental area (VTA)^14^. Stroke is known to disrupt both inhibitory and dopaminergic systems^6,15,16^, complicating the interpretation of prior findings. Therefore, accurate investigation of LTP after stroke requires an experimental model where these modulatory inputs are preserved. In this study, we developed an *in vivo* preparation that maintains endogenous inhibitory tone and neuromodulatory input, enabling induction of LTP through convergent VTA–M1 activation without pharmacological intervention. This approach allowed a precise assessment of LTP in the injured brain.

## Material and Methods

### Ethical approval

Experiments were conducted in accordance with the European Council Directive of 24 November 1986 (86/609/EEC), with the Swiss regulations and approved by the Federal Veterinary Office of Switzerland (licenses #ZH071/2020 and ZH005/2023). All efforts were made to minimize the number of animals used and their suffering. Protocols prepared for this study were not pre-registered. Data created for the study are available in a persistent repository.

### Animals and study design

A total of 84 adult Sprague-Dawley rats (9-11 weeks at the time of the experiment, RjHan:SD, Janvier, France) of both sexes were used for the study. Rats were pair-housed in T2000 IVC cages under an inverted 12-hour light/dark cycle. Access to food and water were provided *ad libitum*.

### Experiment 1: Development of an *in vivo* model of LTP induction in the motor cortex

In this experiment, we developed an effective stimulation protocol for LTP induction in M1 allowing the investigation of LTP in near-physiological conditions. Forty-four rats (21 females) were randomly allocated into 3 experimental groups (M1 only, VTA only, paired VTA-M1) using block randomization. Twenty rats were excluded because of technical failures e.g., broken electrodes (n = 8), premature death (n = 11) or uncontrollable bleeding (n = 1). No sample size analysis was performed for this set of experiments because the goal was to demonstrate the feasibility of inducing LTP in M1 without pharmacological intervention. The data presented here were drawn from a total sample size of 24 animals.

### Experiment 2: LTP induction after stroke

In this experiment, we assessed LTP induction in the peri-infarct cortex after stroke in 36 rats (14 females). In each cage, one rat was randomly assigned to receive either sham surgery or a focal ischemic lesion. Each cage was then randomly assigned to a post-surgical time point (one-week or two-weeks).

#### Sample size calculations

We based our sample size on the effect size observed in experiment 1 in the only group showing LTP: the paired VTA-M1 group (within-group comparison, f = 0.75). To adjust for potential overestimation of effect size due to lack of blinding, a more conservative effect size of f = 0.5 was used for the power analysis^17^. Using G*Power 3.1.9.7^18^, we determined that a sample size of 15 animals (5/group) was enough to detect a within-subject effect size f = 0.5 for a repeated-measures ANOVA with a within-subject factor (2 time points: pre vs post LTP induction) and a between-subjects factor (3 groups: sham, one-week post-stroke, two-weeks post-stroke). Because the recording electrode had to be placed adjacent to the ischemic lesion, blinding of the experimenter was not feasible.

Of the 36 rats, 21 had to be excluded: dies before experiment concluded (n = 9), showed no elicitable response (n = 4), unstable baseline or DC shift that could not be corrected (n = 7) or misplacement of the VTA stimulation (n = 1). The data presented here are drawn from a total sample size of n = 15.

For both sets of experiments, synaptic plasticity was defined as change in the amplitude of evoked field postsynaptic potentials (fPSPs) of > 10% lasting for at least 30 minutes following the final stimulation. Data were analyzed offline independently from experimental group allocation, allowing for blinded extraction and analysis. All final data analyses were conducted blind to group assignment. Exclusion criteria were established *a priori*. The exclusion rate between experiment 1 and experiment 2 were not statistically different (χ^2^ = 0.85, df = 1, p = 0.357).

### Experimental procedure

The rats’ body temperature was maintained at 36-37°C with a warming pad controlled with a rectal probe. Eye ointment (Vitamin A, Bauch and Lomb, Switzerland) was applied to avoid cornea dehydration. Lidocaine (2%, Streuli Pharma AG, Switzerland) diluted 5 times in sterile saline (Braun Medical AG, Germany) was infused under the scalp before opening.

#### Stroke/sham stroke induction

Rats involved in experiment 2 were subjected to a stroke or sham stroke surgery in the left hemisphere. The heart rate, respiration rate and body temperature were continuously monitored during the procedure (PhysioSuite, Kent Scientific, USA). Oxygen was provided during the surgery through a nose piece (929 B, Kopf Instruments, USA). The adequate depth of anaesthesia was evaluated by regularly assessing the absence of a nociceptive reflex to a toe pinch as well as the absence of reaction to a light ocular touch. Additional injection of anaesthetics was given as needed.

Rats were anesthetized with a subcutaneous (s.c.) injection of a triple mix of fentanyl (0.005 mg/kg, Sintetica, Switzerland), midazolam (2 mg/kg, Dormicum, Roche, Switzerland) and medetomidine (0.15 mg/kg, Domitor, Orion Pharma, Finland). Meloxicam (1 mg/kg, Metacam 2mg/ml, Boehringer, Switzerland) was injected s.c. before and 24 hours after surgery to prevent surgical pain. The head of the rat was shaved and secured in a stereotactic frame (1430, Kopf Instruments, USA) before being cleaned with alcohol and betadine. An incision was performed along the midline axis, and the conjunctive tissues were cleaned to expose the skull. The bregma and lambda were aligned along the horizontal axis, and a small craniotomy was drilled over M1 using a dental drill (Foredom, USA) at the following coordinates relative to bregma: 2.3 mm anterior and 3 mm lateral. A stroke was induced by the injection of 1 microliter of human and porcine endothelin-1 (ET-1; 400 micromolar in sterile PBS, Merck-Millipore, USA) 1.5 mm under the pial surface. The injection was performed at a speed of 1 microliter/min using a Hamilton syringe (84851, Hamilton Company, USA) mounted on a motorized stereotactic injector (QSI, Stoelting Co., USA). The syringe was left in place for about 3 minutes before being slowly withdrawn to prevent backflow. Sham rats were subjected to the same procedure except that 1 microliter of sterile PBS was injected instead of ET-1 in PBS. The craniotomy was then closed with dental cement, and the scalp incision was sutured and covered with betadine. The rats received an injection of warm Ringer Lactate solution (2 ml, s.c., Braun Medical AG, Germany). The anaesthetics were antagonized with an intraperitoneal (i.p.) injection of a triple mix of buprenorphine (0.05 mg/kg, Temgesic, Indivior, Switzerland), atipamezole (0.75 mg/kg, Revertor, Virbac, France), and flumazenil (0.2 mg/kg, Anexate, Cito Pharma Service GmbH, Switzerland). The rats were allowed to recover in a clean cage placed over a heating pad before returning to their home cage.

#### Electrophysiological recordings

Coordinates represent millimeters relative to bregma. Evoked field potentials (fPSPs) were recorded in M1 using electrodes pulled from glass capillaries (BF-150-86-10, Sutter Instruments, USA) using a horizontal puller (Model P97 Flaming Brown, Sutter Instruments, USA) and filled with 0.4 M NaCl. Analog signal was amplified 10 000 times and bandwidth filtered (0.1Hz-1kHz) using a Multiclamp 700B (Molecular Devices, USA) before being digitized and saved on a computer using a data acquisition board (National Instruments, USA) and custom-made software in LabVIEW 2019 x64 (National Instruments, USA). The synaptic strength was assessed by measuring the peak amplitudes of fPSPs offline using LabVIEW 2019 x64.

#### Surgical procedure

Rats were anesthetized with an i.p. injection of urethane (1.75 mg/kg, Sigma Aldrich, USA) and received a s.c. injection of meloxicam (1 mg/kg, Metacam 2mg/ml, Boehringer, Switzerland). The head was shaved and placed in a stereotactic frame (1430, Kopf Instruments, USA). A midline incision was performed, and the skull was cleaned. The bregma and the lambda were aligned in the horizontal axis before performing a craniotomy centred over the left M1 (2.3 mm anterior, 3 mm lateral) and another one at coordinates 6.23 mm posterior, 3.05 mm lateral to reach the left VTA with an angle of 20°. The border of the stroke was identified by the demarcation between the ischemic region showing a white color and the non-ischemic surroundings showing a yellow color. A concentric bipolar stimulation electrode (30205, FHC Inc, USA) was placed in M1 or the peri-infarct cortex in layer II/III 250 µm under the pial surface and a similar stimulation electrode was placed in the VTA 7.45 mm under the pial surface. Finally, a recording electrode was placed in M1 or the peri-infarct area in close vicinity to the stimulation electrode 250 to 300 µm under the pial surface. A reference electrode was placed under the skin of the rat above the cerebellum. To prevent brain swelling during the experiment, the skull was kept moist with artificial cerebrospinal fluid of following composition (in mM): 126 NaCl, 3 KCl, 1.25 NaH_2_PO_4_, 26 NaHCO_3_, 1 MgSO_4_, 2 CaCl_2_ and 10 glucose. Recordings started about 20 minutes after the electrodes were in place.

#### LTP induction

The following methods were adapted from a classical paradigm to induce LTP in M1 in slice preparation^3,9,12^. fPSPs were evoked using current paired pulses in M1 (0.2 ms duration, 50 ms apart) every 30 seconds. First, a threshold stimulation intensity was defined as the stimulation intensity necessary to evoke an fPSP amplitude of approximately 0.2 mV. Then, synaptic strength was assessed for a range of stimulation intensities using multiples of threshold intensity to build an input-output (IO) curve (1 to 5x threshold intensity in 0.5x increment). For each stimulation intensity, three fPSPs were recorded. The subsequent recordings were performed using a stimulation intensity necessary to evoke an fPSP response amplitude of approximately 50% the maximum amplitude recorded during the IO curve. After a 20 minutes baseline, induction of LTP was attempted 2-3 times separated by a 20–30-minute recording period in naïve rats following 3 different protocols detailed below. After the last attempt, fPSPs were recorded for 30 minutes (Fig. 1A).

**Fig. 1:**
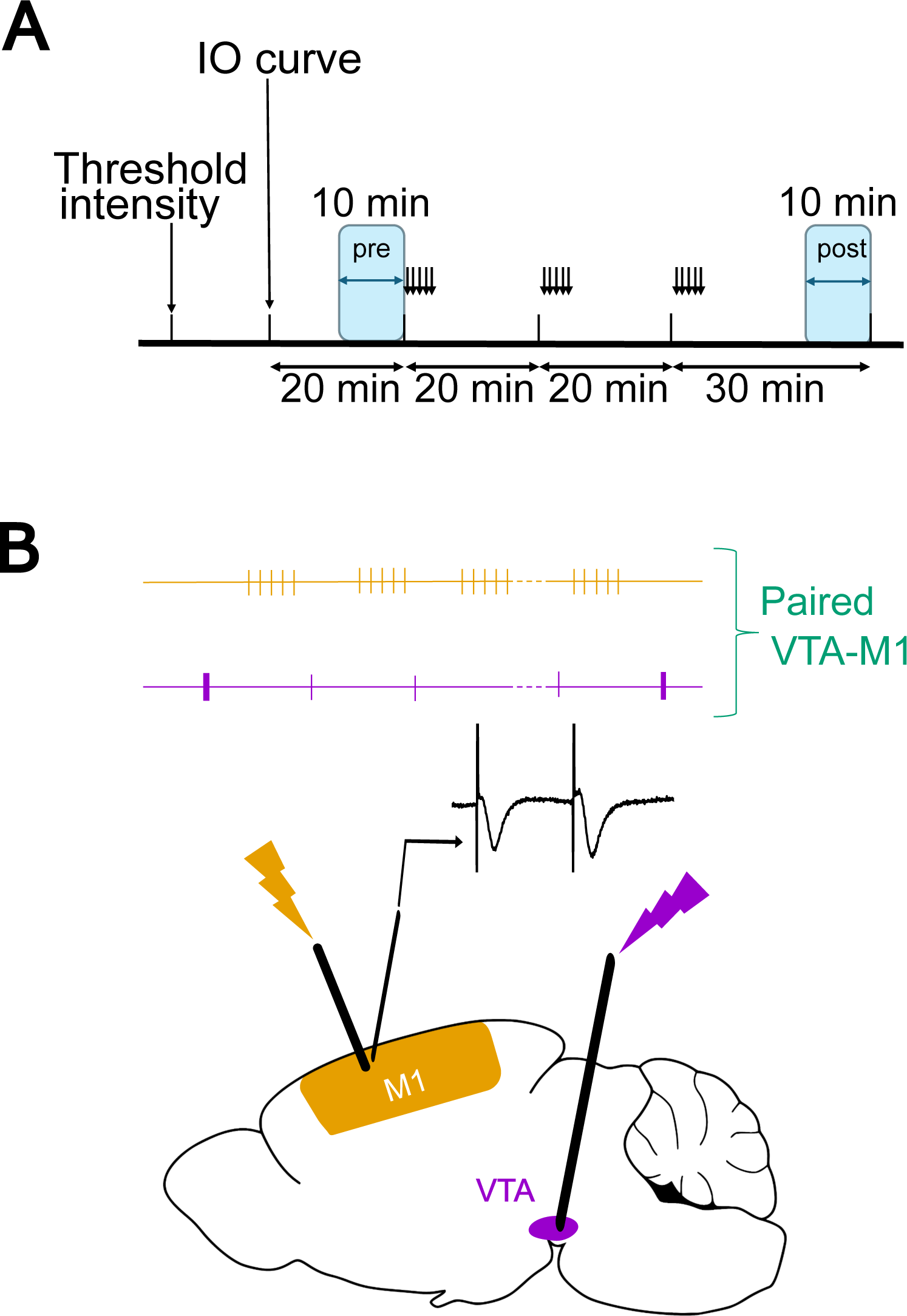
Schematic representation of the experiments. A – Experimental timeline. After determination of the threshold intensity, an IO curve was recorded to find the stimulation intensity evoking the maximal fPSPs amplitude. Then, the stimulation intensity to evoke fPSPs of half the maximum amplitude was used to record fPSPs in blocks of 20 minutes interspersed with attempts to induce LTP (black arrows). Changes in fPSPs amplitudes were evaluated by comparing the averaged response during the last 10 minutes recording (light blue frame, “post”) to the average baseline amplitude recorded during 10 minutes before the first attempt (light blue frame, “pre”). B – Schematic representation of the experimental setup. Concentric bipolar stimulation electrodes were implanted in M1 and the VTA and a glass recording electrode in M1 to evoke M1 responses to M1 paired-pulse stimulations. Three different protocols were used to attempt inducing LTP: TBS in M1 (yellow), a tonic (4 Hz, thin purple ticks) stimulation of the VTA flanked by two phasic (20 Hz, thick purple ticks) or a combination of both protocols (green).

Induction of LTP was attempted using 3 different stimulation protocols in naïve rats: either theta burst stimulation (TBS) in the M1 only group, a phasic-tonic-phasic stimulation in the VTA only group, or simultaneous stimulation of VTA and M1 in the paired VTA-M1 group. In sham and stroke rats, induction of LTP was only attempted with simultaneous VTA-M1 stimulation (Fig. 1B). The TBS protocol in M1 consisted of 10 trains of 5Hz stimuli, each composed of 4 pulses delivered at 100 Hz, repeated 5 times every 10 seconds (Fig. 1B, yellow trace). VTA stimulation consisted of a 2 second stimulation at 4 Hz, to mimic tonic activation of the VTA, flanked by two periods of 2 seconds during which the VTA was stimulated at 20 Hz, to mimic phasic activation of the VTA (Fig. 1B, purple trace). During the paired M1-VTA protocol, the M1 TBS protocol and the VTA stimulation protocol were synchronized so that each repetition of the TBS protocol was preceded and followed by a 20 Hz stimulation period (Fig. 1B, yellow and purple trace). Pulses were 200 µseconds long. To induce LTP, the stimulation intensity in M1 was set at twice the stimulation intensity used for baseline recording^3,9,12^. However, no pharmacological blockade of the GABAergic receptors was used in this study.

At the end of the recordings, the rats were decapitated under urethane anaesthesia, the brain was removed and immersed into 4% paraformaldehyde in 0.1M PBS for anatomical verification of the stimulation electrode placement in the VTA and the stroke placement using a classical Nissl staining procedure.

#### Assessment of motor function

Before sham stroke/stroke induction and before electrophysiological recordings, rats’ motor function was assessed using the cylinder test^19^. The rats were placed in a Plexiglas cylinder (20 cm diameter, 35 cm high). The number of paw placements on the vertical cylinder wall was counted for the left and the right paw separately for 3 minutes

### Data analysis

For electrophysiological data, individual IO curves were constructed by averaging the three fPSP peak amplitudes for each stimulation intensity. The baseline and post-induction synaptic strengths were computed by averaging the last 20 fPSPs recorded before the first attempt to induce plasticity and the last 20 fPSPs recorded 30 minutes after the last attempt to induce plasticity, respectively (10 minutes recordings, Fig1. A). Paired-pulse ratio (PPR) was calculated as the ratio of the second peak amplitude to the first peak amplitude.

For behavioral data, the ratio between the right paw and the total number of touches was then calculated.

### Statistical analysis

Statistical analyses were performed with R 4.3.2 in RStudio 2023.09.1 using rstatix, lmerTest and emmeans packages^20–22^. To test for a difference in exclusion rate between experiment 1 and 2, a chi-square test was used. For all remaining statistical tests, normality and homogeneity of variance were tested using a QQ plot and a Scale-Location plot, respectively to inform on the use of a parametric or non-parametric test.

In experiment 1, the effects of the three stimulation protocols on synaptic strength were assessed using a type III ANOVA on a linear mixed model including GROUP (between-subject factor, 3 levels: M1 only, VTA only, paired VTA-M1), TIME POINTS (within-subject factor, 2 levels: pre, post) and their interaction as fixed factors and rats as a random factor. Short-term plasticity was assessed using the paired-pulse ratio (PPR). To test whether the PPR is different than µ = 1, a one-sample t-test was used. The similarity of baseline conditions across groups was assessed using a Welch’s one way ANOVA on baseline fPSPs amplitudes and baseline stimulation intensities and a type III ANOVA on a linear model on the IO curves including STIMULATION INTENSITY (within-subject factor, 10 levels: 1 to 5x threshold intensity in 0.5x increment), GROUP (between-subject factor, 3 levels) and their interaction as fixed factors and rats as random factor.

In experiment 2, the effect of a stroke on synaptic strength was assessed using a type III ANOVA on a linear mixed model including GROUP (between-subject factor, 3 levels: Sham, one-week post-stroke, two-weeks post-stroke), TIME POINTS (within-subject factor, 2 levels: pre, post) and their interaction as fixed factors and rats as a random factor. To test for the effect of stroke on short-term plasticity, a Kruskall Wallis test on the PPR was used. To assess motor function in sham and stroke rats, we used a type III ANOVA on linear models for the number of contacts with the cylinder and on the paw use asymmetry. For each outcome, the linear model including the GROUP (between-subject factor, 3 levels) and the TIME POINTS (within-subject factor, 2 levels) as fixed factors and rats as random factor. The effect of stroke on baseline synaptic transmission was assessed using a type III ANOVA on a linear model on the IO curves including STIMULATION INTENSITY (within-subject factor, 10 levels: 1 to 5x threshold intensity in 0.5x increment), GROUP (between-subject factor, 3 levels) and their interaction as fixed factors and rats as random factor, a Welch’s one way ANOVA on baseline amplitude and a Kruskall Wallis test on baseline stimulation intensities.

The change in peak amplitudes was used to analyse LTP induction rather than percentage of baseline to prevent misleading results^23^. To increase clarity, results are expressed as peak amplitudes and percentages of baseline. Pairwise t-test with Kenward-Roger degrees of freedom on the marginal means with Holm correction were performed for post-hoc comparisons after the ANOVAs. The Kenward-Roger estimation of degrees of freedom showed good accuracy for small samples^24^. Data are presented as mean ± SD unless otherwise specified. Significance was considered for p < 0.05.

## Results

### Simultaneous VTA-M1 activation is required to induce LTP in M1 *in vivo*

To investigate synaptic plasticity under near *in vivo* conditions, we placed concentric stimulation and recording electrodes in layers II/III of M1 and a concentric stimulation electrode in VTA in urethane-anesthetized rats. Paired pulse stimulation of M1 (0.2 ms duration, ISI = 50 ms, frequency = 0.033Hz) reliably evoked negative-ongoing responses peaking at 8.63 ± 1.08 ms. The PPR was significantly greater than 1 (1.14 ± 0.17, t_23_ = 4.06, p = 0.0005, n = 24) consistent with short-term facilitation.

To induce LTP, we compared three stimulation conditions: M1-only (TBS), VTA-only (phasic-tonic stimulation), and simultaneously paired VTA-M1 stimulation (Fig. 1B). Only the paired protocol resulted in significant potentiation of fPSP amplitudes to 114.35 ± 12.47% of baseline (GROUP x TIMEPOINT: F_2,24_ = 4.76, p = 0.018_;_ pre vs post: 0.56 ± 0.15 mV vs 0.64 ± 0.15 mV, t_27.4_ = 3.36, p = 0.007, n = 8; Fig. 2A, B). Neither M1-only, nor VTA-only stimulation altered synaptic strength (pre vs post, M1 only: 0.69 ± 0.26 vs 0.68 ± 0.26, t_27.4_ = 0.29, p = 1, 99.25 ± 7.22% of baseline; VTA only: 0.64 ± 0.15 vs 0.64 ± 0.12, t_27.4_ = 0.06, p = 1, 101.12 ± 12.08% of baseline; n = 8 for both groups; Fig. 2A,B).

**Fig. 2:**
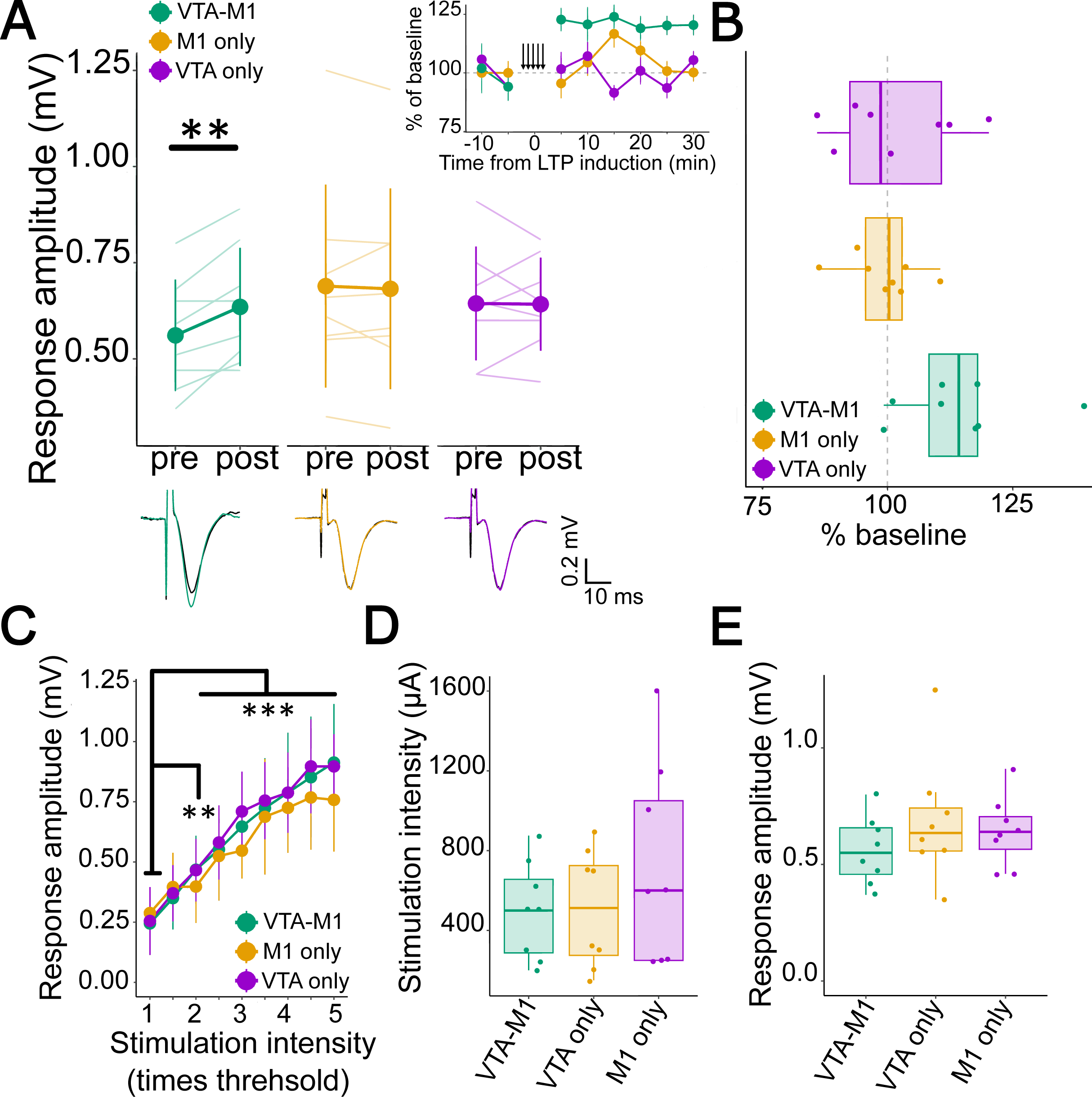
Induction of LTP in the motor cortex necessitates simultaneous VTA and M1 activation. A – LTP of fPSP amplitudes was observed only with paired VTA–M1 stimulation. fPSP amplitudes were significantly increased post-stimulation (post) compared to baseline (pre) in the paired group, but neither M1 nor VTA stimulation alone. Representative traces before (black) and after (colored) LTP induction are displayed below the graph for each experimental group. Solid lines and points represent group means ± SD. Transparent lines show individual data. Inset shows a representative time course of LTP induction in the peri-infarct cortex for each experimental group. Data points represent 5-minute averages (mean ± SEM) of the peak fPSP amplitudes at baseline and after the final induction attempt (indicated by the vertical arrows). B – Increase in fPSP amplitudes shown in A represented in percentage of baseline. C – Input-output (IO) curves. Increasing stimulation intensity significantly increased fPSP amplitudes. No group differences were detected. **: p <0.01; ***:p < 0.001 compared to 1x threshold intensity. D – Baseline stimulation intensities did not differ between groups. E – Baseline response amplitude was also comparable between groups. A-E – Purple = VTA only; Yellow = M1 only, Green = Paired VTA-M1; n = 8 per group. A, B inset: A (inset), B: The grey dashed line indicates normalized baseline. D, E: Boxplots show median and interquartile range. Points represent individual animal data.

These results were unlikely due to a difference in synaptic strength during baseline or to different experimental conditions between groups. IO curves showed the expected amplitude increase with rising stimulation intensity, with no significant group differences (STIMULATION INTENSITY: F_8,192_ = 110.22, p < 2e^-16^, 1.5x threshold intensity vs 1x threshold intensity, t_219_ = 3.59, p = 0.004, all other multiple thresholds vs 1x threshold: ps < 0.0001; GROUP: F_2,24_ = 0.56, p = 0.58; Fig. 2C). Baseline stimulation intensities (M1-only: 509.38 ± 295.8 µA, VTA-only: 718.75 ± 503.51 µA, paired VTA-M1: 500 ± 242.38 µA, F_2,13.19_ = 0.61, p = 0.56; Fig. 2D) and initial fPSP amplitudes (M1-only: 0.69 ± 0.26, VTA-only: 0.64 ± 0.15, paired VTA-M1: 0.56 ± 0.15, F_2,13.4_ = 0.96, p = 0.41; Fig. 2E) were also equivalent. These data demonstrate that LTP can be induced in M1 of anesthetized rats only when M1 stimulation is paired with concurrent VTA input, reflecting the necessity of intact neuromodulatory drive for synaptic plasticity under quasi-natural conditions.

### Stroke impaired motor function

To study the capacity of the peri-infarct cortex to express LTP, we applied the validated VTA-M1 protocol in rats with ET-1-induced ischemia in the M1 forelimb region (Fig. 3A).

**Fig. 3:**
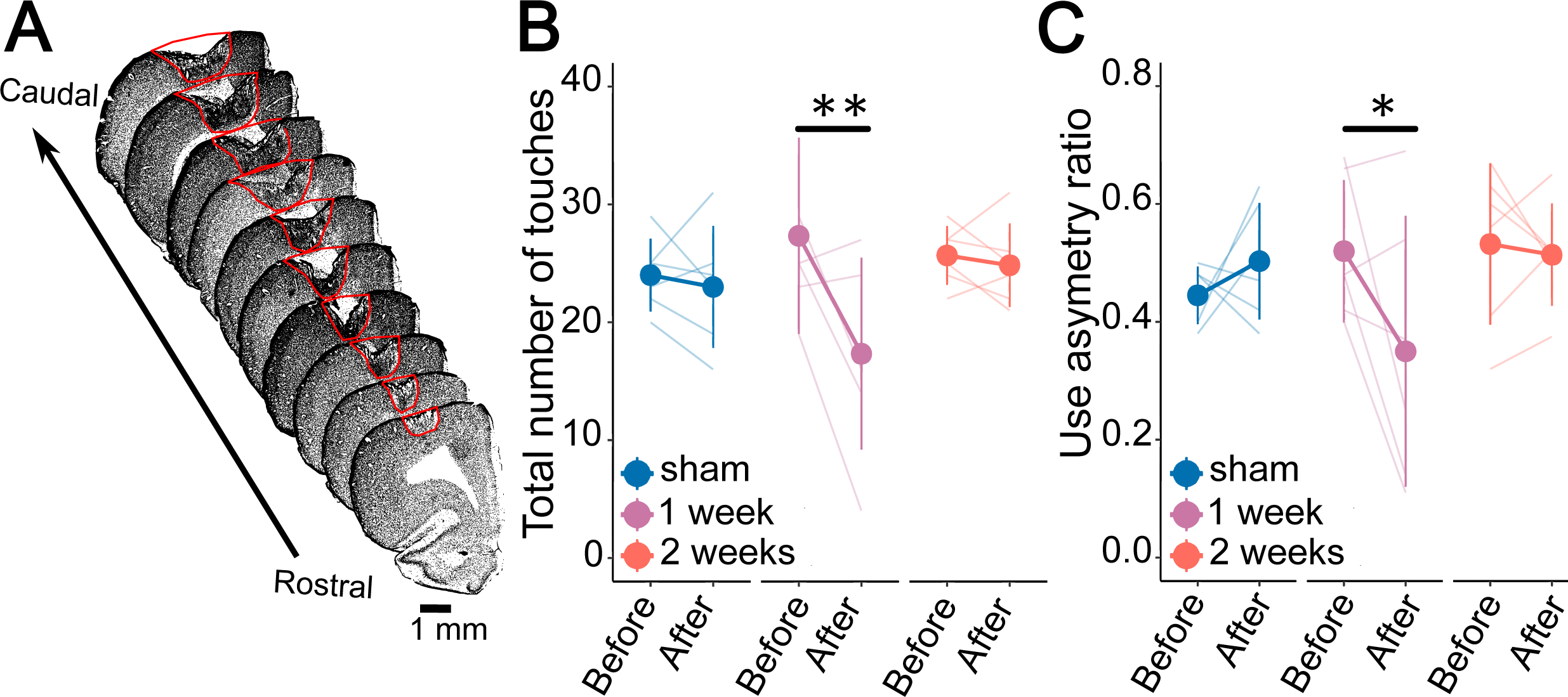
Stroke impaired motor function. A – Representative photomicrographs of a stroke outlined in red. B –Exploratory behavior in the cylinder test before and after stroke. Exploratory behavior was significantly reduced one week after stroke, as measured by the number of wall contacts in the cylinder test. No change was observed in sham-operated rats or at two weeks post-stroke. C – Paw use asymmetry in the cylinder test before and after stroke. Contralesional forelimb use was also reduced one week after stroke, with recovery by two weeks. Sham rats showed no significant change. *: p < 0.05, **: p < 0.01 before vs after stroke. Green = sham, blue = one week after stroke, yellow = two weeks after stroke; n = 6 per group.

Stroke produced transient, behaviorally relevant deficits: exploratory behavior (cylinder contacts, TIMEPOINT: F_1,15_ = 6.51, p = 0.02; GROUP X TIMEPOINT: F_2,15_ = 3.84, p = 0.045) was reduced in rats tested at one-week (27.3 ± 8.3 vs 17.3 ± 8.1, t_15_ = -3.74, *p* = 0.002, n = 6) but not in rats tested at two-weeks post-stroke (25.67 ± 2.5 vs 24.83 ± 3.5, t_15_ = 0.31, *p* = 0.76, *n* = 6; Fig. 3B). Contralesional paw use (GROUP X TIMEPOINT: F_2,18_ = 3.82, p = 0.041) declined at one-week (0.52 ± 0.12 vs 0.35 ± 0.23, t_21.6_ = 2.61, p = 0.02, n = 6) and did not show a difference at two-weeks post-stroke (0.53 ± 0.14 vs 0.51 ± 0.09, t_21.6_ = 0.27, p = 0.79, n = 6; Fig. 3C). As expected, sham surgery neither modified the number of contacts with the cylinder (before vs after: 24 ± 3.1 vs 23 ± 5.18, t_15_ = 0.37, p = 0.71, n = 6; Fig. 3B) nor the paw use asymmetry (before vs after: 0.45 ± 0. 05 vs 0.5 ± 0.1, t_21.6_ = 0.9, p = 0.38, n = 6; Fig. 3C).

### Synaptic plasticity is impaired in the peri-infarct region after stroke

Applying the paired VTA-M1 stimulation protocol established in the healthy condition one- and two-weeks post-stroke differentially affected the fPSP amplitudes between the groups (GROUP x TIMEPOINT: F_2,12_ = 11.35, p = 0.002, Fig. 4A, B). In the sham group, fPSPs amplitude were significantly enhanced (before vs after: 1.00 ± 0.35 mV vs 1.42 ± 0.3 mV, t_18.8_ = 3.71, p = 0.005, 149 ± 41.21% of baseline, n = 4, Fig. 4A, B). In the stroke group, however, fPSP amplitude was significantly depressed at one-week post-stroke (before vs after: 0.65 ± 0.42 mV vs 0.40 ± 0.18 mV, t_18.8_ = 2.63, p = 0.03, 76.27 ± 36.51% of baseline, n = 6, Fig. 4A,B), and at two-weeks post-stroke, although it was no longer significantly different from baseline (before vs after: 0.56 ± 0.31 mV vs 0.40 ± 0.21, t_18.8_ = 1.58, 73.74 ± 16.94% of baseline, p = 0.13, n = 5, Fig. 4A,B). These results indicate that induction of LTP is supressed in the peri-infarct area.

**Fig. 4:**
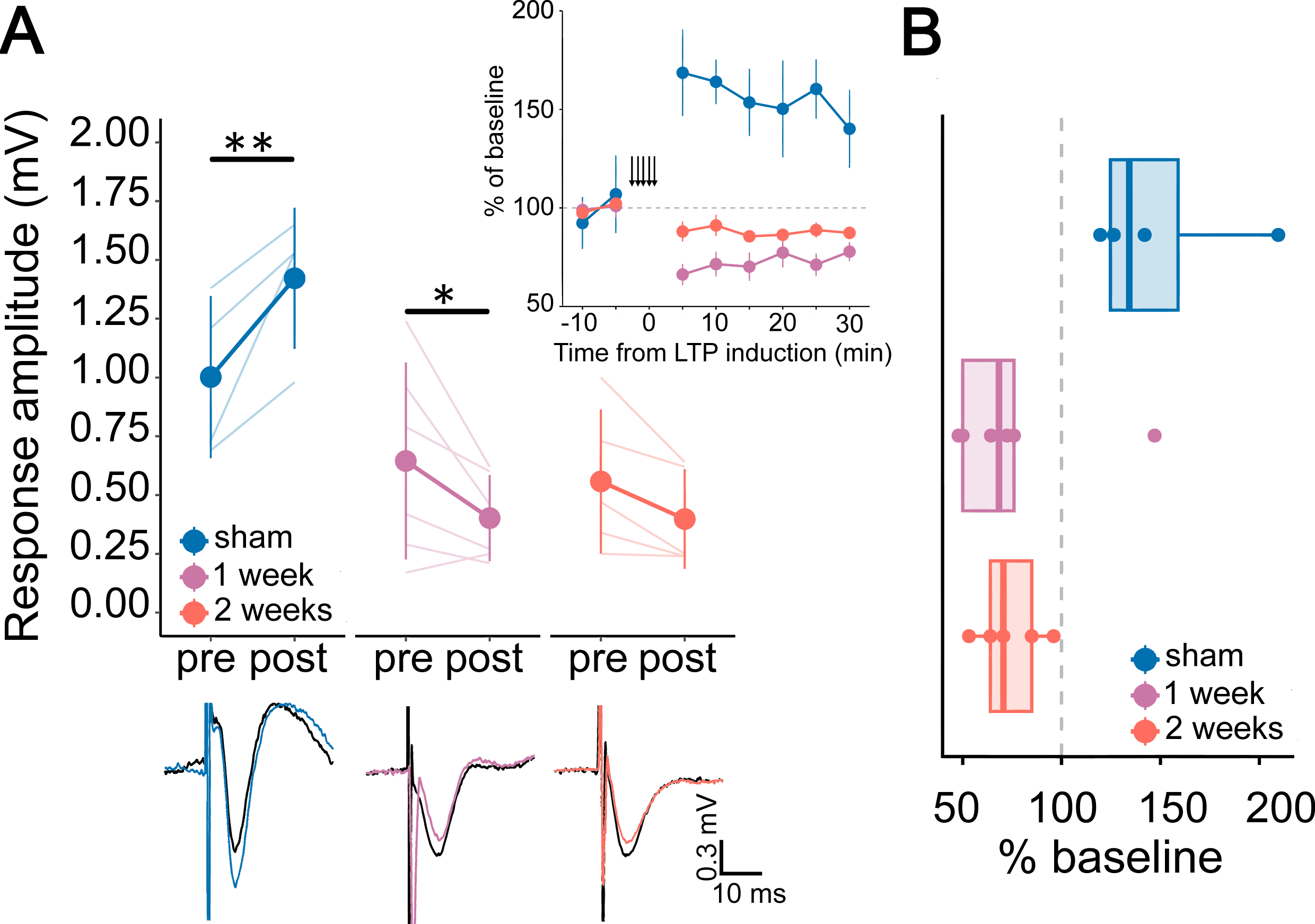
Impairment of synaptic plasticity in the peri-infarct area after stroke. A -– Effects of paired VTA-M1 stimulation on fPSP amplitudes in the peri-infarct area. LTP was observed in sham rats but not after stroke. Response amplitudes significantly increased post-stimulation in the sham group but remained unchanged or decreased 1 week or 2 weeks after stroke. Representative traces before (black, pre) and after (colored, post) LTP induction are displayed beneath the graph. Solid lines/points show group mean and ± SD; transparent lines represent individual animal data. Inset shows a representative time course of LTP induction in the peri-infarct cortex for each experimental group. Data points represent 5-minute averages (mean ± SEM) of the peak fPSP amplitudes at baseline and after the final induction attempt (indicated by the vertical arrows). B-Increase fPSP amplitudes shown in A represented in percentage of baseline. A-F : Blue = sham (*n* = 4); Light purple = one-week post-stroke (*n* = 6); Light orange = two-weeks post-stroke (*n* = 5). In A (inset) and B, the grey dashed line indicates normalized baseline. In D and E, boxplots show median and interquartile range; individual data points are overlaid.

**Fig. 5:**
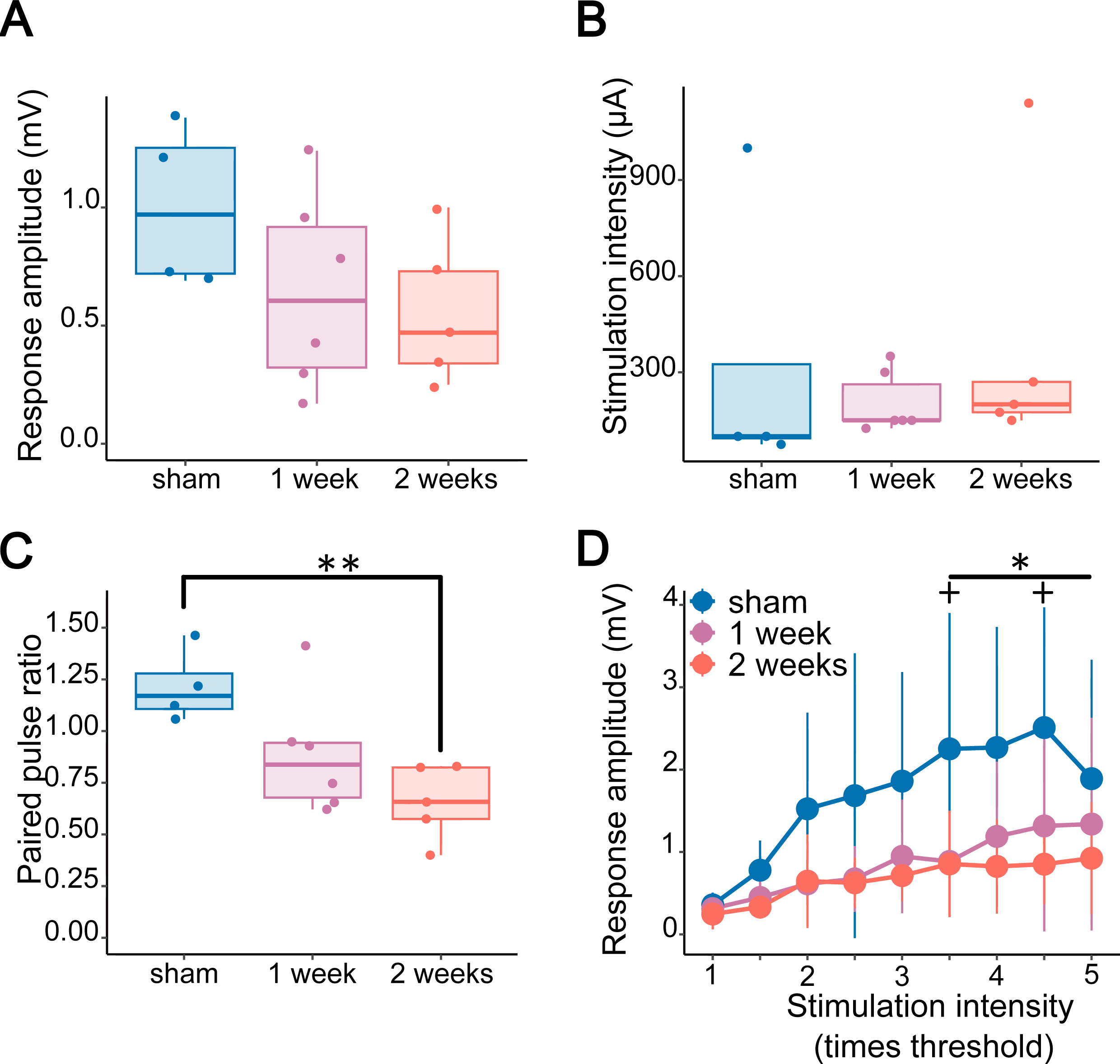
Impairment of synaptic transmission after stroke. A – Response amplitudes at baseline. Baseline fPSP amplitudes did not differ between groups. B – Stimulation intensity at baseline. Baseline stimulation intensities were also comparable across groups. C – Short-term plasticity. Paired-pulse ratio (PPR) decreased progressively post-stroke, indicating a shift from short-term facilitation (PPR > 1) to depression (PPR < 1), with a significant reduction two weeks after stroke compared to sham. D – IO curves revealed impaired synaptic gain post-stroke. Higher stimulation intensities elicited significantly smaller fPSPs one week and two weeks after stroke compared to sham. +: sham against one-week, *: sham against two-weeks

### Stroke disrupts synaptic transmission and short-term plasticity

We then tested whether baseline synaptic transmission was impaired after stroke. Baseline fPSP amplitudes (sham: 1 ± 0.35 mV, n = 4; one-week: 0.65 ± 0.42 mV, n = 6; two-weeks: 0.56 ± 0.31 mV, n = 5; F_2,7.5_ = 1.95, p = 0.21, Fig. 4C) and stimulation intensities (sham: 318.75 ± 454.32 µA, n = 4; one-week: 204.17 ± 95.42 µA, n = 6; two-weeks: 387 ± 423.31 µA, n = 5; H_2_ = 3.19, p = 0.2, Fig. 4D) did not differ across groups. However, PPR decreased progressively post-stroke with a significant difference between the stroke and sham control group at two-weeks post-stroke (F_2,7.78_ = 9.98, p = 0.007; sham vs one-week: 1.22 ± 0.18 vs 0.89 ± 0.29, p = 0.05; sham vs two-weeks: 1.22 ± 0.18 vs 0.66 ± 0.18, p = 0.008, Fig. 4E). In addition, the PPR at two-weeks post-stroke was significantly below 1, indicating a shift from facilitation to short-term depression (t_4_ = 4.24, p = 0.01, Fig. 4E).

IO curves revealed impaired synaptic recruitment after stroke (GROUP X INTENSITY: F_16,120_ = 2.43, p = 0.003, Fig. 4F). Higher stimulation intensities evoked lower fPSP amplitudes one- and two-weeks post-stroke compared to sham (p ranging from p = 0.04 to p = 0.002 for 3.5x threshold and above) with a marginal non-significance at one-week post-stroke for 4x threshold (p = 0.06) and 5x threshold (p = 0.05), indicating that stroke gradually reduced the range of responses in the peri-infarct area for stronger inputs as for PPR. Consequently, the peri-infarct area was less responsive to increasing stimulation intensities compared to sham. Although 2x threshold intensity was enough to induce a significantly larger fPSPs in the sham group (1x threshold vs 2x threshold: t_150_ = 3.4, p = 0.002, n = 4), a difference was only detectable at 4x threshold intensity one-week post-stroke (1x threshold vs 4x threshold: t_150_ = 3.1, p = 0.013, n = 6) and no difference was detected even at 5x threshold intensity two-weeks post-stroke (1x threshold vs 5x threshold: t_150_ = 2.21, p = 0.23, n = 5). In conclusion, in our model, stroke led to a durable impairment of both synaptic transmission and long-term plasticity in the peri-infarct cortex, likely reflecting a degraded capacity for synaptic potentiation and reduced circuit responsiveness.

## Discussion

Although increasingly debated, many still assume that a time-limited window of heightened plasticity following stroke supports behavioral recovery through mechanisms akin to those underlying motor learning^25–28^. Among these, LTP has been proposed as a central mechanism, based primarily on findings showing high structural remodelling and overexpression of neurotrophic factors involved in healthy motor learning^1,27,29^. Although earlier work in slice electrophysiology interrogated the capacity of the peri-infarct area to express LTP at time points suitable for recovery^9–11^, this has rarely been tested *in vivo* under conditions that preserve intact circuit dynamics. In fact, methodological constraints in previous studies may have obscured critical inhibitory mechanisms or long-range modulatory inputs that shape recovery. In this study, we developed a minimally perturbed *in vivo* preparation enabling the study of the capacity of the peri-infarct cortex to express LTP without pharmacological confounds. We found a markedly reduced capacity for LTP and synaptic transmission one and two weeks after stroke despite good motor recovery.

Our findings add to a growing body of evidence suggesting that increased structural remodelling in the peri-infarct cortex may not necessarily equate to increased functional plasticity^30–32^. While prior work has reported elevated dendritic spine turnover and axonal sprouting in this region during early recovery^26,33,34^, our results imply that such remodelling may occur in a physiologically constrained, or even dysfunctional, environment. Consistent with this view, similar impairments in LTP have been observed in the hippocampus following ischemia^10,11^, even in the absence of local structural damage.

The idea that stroke-induced plasticity enhances learning capacity has been supported by molecular parallels between spontaneous recovery and motor learning^27,29^. This has led to the suggestion that early post-stroke brains are more responsive to training because they are “better learners”. However, recent behavioral evidence—including work from our group— has called this into question. For example, while performance in error-based motor learning tasks remains intact early after stroke, reinforcement-based learning appears selectively impaired, even when lesion location does not directly involve the reward system^35^. This is consistent with broader evidence of acute dysfunction in reward-related networks following stroke and may help explain unexpected cognitive and motivational deficits observed even after small, subcortical lesions^36–38^.

Our results suggest that LTP suppression after stroke is not simply due to inadequate stimulation but reflects a deeper disruption in synaptic integration. The shift from short-term facilitation to depression, along with reduced synaptic responses at higher stimulation intensities, points to increased inhibitory tone or impaired dendritic excitability as likely contributors. Prior computational and experimental work has shown that increased tonic inhibition can reduce synaptic gain and flatten IO curves^39,40^, potentially rendering the peri-infarct cortex unresponsive to potentiation protocols. Our results are consistent with previous studies in rats demonstrating a long-lasting reduction in sensory response amplitude in the sensory cortex after stroke^41^ and could provide a mechanism for the lack of neural responsiveness during increasing cognitive task difficulty in stroke patients who recovered from minor strokes^42^. Noteworthy, although cognitive recovery is paralleled by increased interhemispheric connectivity compared to healthy controls, the restoration of neural dynamics closer to those seen in healthy controls appears key to the maintenance of the cognitive function^43^. This emphasizes the idea of the existence of an optimal functional brain state^44^ with an upper and lower limit beyond which cardinal neural features, such as LTP, cannot be supported anymore. The restoration of this critical state might be preponderant for motor recovery.

Methodologically, our approach addresses an important gap. Earlier studies investigating LTP after stroke used slice electrophysiology^9–11^. Although this classical preparation to study LTP is powerful to precisely investigate the mechanisms of LTP, it may fail to account for long-range influences and brain changes caused by a stroke^6,15^. In contrast, we developed a minimally perturbed *in vivo* protocol that permits LTP induction through converging VTA–M1 activation without pharmacological intervention. Our results therefore offer a more accurate reflection of synaptic plasticity potential in the injured brain. Indeed, the inability to induce LTP in stroke rats occurred despite the use of dopaminergic stimulation parameters previously shown to enhance cortical firing and dopamine release under anesthesia^45,46^.

Overall, our data indicate profound functional alterations in synaptic transmission after stroke associated with impairment of LTP in the peri-infarct cortex despite significant motor recovery. The dramatic structural changes happening in the peri-infarct area are often interpreted as the marker for a hyperplastic milieu that could be leveraged to augment recovery through learning-induced LTP. Although structural plasticity and LTP are necessary to mediate motor skill learning, they are independent mechanisms^47^, with LTP stabilizing synapses relevant for the new skill to acquire^48,49^. Therefore, our data suggest that structural hyperplasticity of the peri-infarct area early after stroke cannot support learning-induced improvement; instead, it may reflect a destabilized environment that cannot support LTP induction^30,50^. They also provide a biological basis to more recent evidence showing deficit in specific forms of learning after stroke^35,36^. These results align with an emerging framework in which recovery is mediated by the restoration of networks dynamics rather than local synaptic reconnection^43,44^.

## Authors contribution: CRediT

CV: Conceptualization, Methodology, Software, Formal analysis, Investigation, Data Curation, Writing – Original Draft

MB: Writing – Review and Editing WJ: Writing – Review and Editing

AL: Funding acquisition, Resources, Conceptualization, Supervision, Writing – Review & Editing

## References

1. Biernaskie J, Chernenko G, Corbett D. Efficacy of rehabilitative experience declines with time after focal ischemic brain injury. J Neurosci. 2004;24:1245–1254.

2. Dromerick AW, Geed S, Barth J, Brady K, Giannetti ML, Mitchell A, Edwardson MA, Tan MT, Zhou Y, Newport EL, et al. Critical Period After Stroke Study (CPASS): A phase II clinical trial testing an optimal time for motor recovery after stroke in humans. Proc Natl Acad Sci U S A. 2021;118:e2026676118.

3. Rioult-Pedotti MS, Friedman D, Hess G, Donoghue JP. Strengthening of horizontal cortical connections following skill learning. Nat Neurosci. 1998;1:230–234.

4. Nabavi S, Fox R, Proulx CD, Lin JY, Tsien RY, Malinow R. Engineering a memory with LTD and LTP. Nature. 2014;511:348–352.

5. Hill TC, Zito K. LTP-induced long-term stabilization of individual nascent dendritic spines. J Neurosci. 2013;33:678–686.

6. Clarkson AN, Huang BS, Macisaac SE, Mody I, Carmichael ST. Reducing excessive GABA-mediated tonic inhibition promotes functional recovery after stroke. Nature. 2010;468:305–309.

7. Lüscher C, Malenka RC. NMDA Receptor-Dependent Long-Term Potentiation and Long-Term Depression (LTP/LTD). Cold Spring Harb Perspect Biol. 2012;4:a005710.

8. Ying S-W, Futter M, Rosenblum K, Webber MJ, Hunt SP, Bliss TVP, Bramham CR. Brain-Derived Neurotrophic Factor Induces Long-Term Potentiation in Intact Adult Hippocampus: Requirement for ERK Activation Coupled to CREB and Upregulation of Arc Synthesis. J Neurosci. 2002;22:1532–1540.

9. Hagemann G, Redecker C, Neumann-Haefelin T, Freund HJ, Witte OW. Increased long-term potentiation in the surround of experimentally induced focal cortical infarction. Ann Neurol. 1998;44:255–258.

10. Li W, Huang R, Shetty RA, Thangthaeng N, Liu R, Chen Z, Sumien N, Rutledge M, Dillon GH, Yuan F, et al. Transient focal cerebral ischemia induces long-term cognitive function deficit in an experimental ischemic stroke model. Neurobiol Dis. 2013;59:18– 25.

11. Orfila JE, Grewal H, Dietz RM, Strnad F, Shimizu T, Moreno M, Schroeder C, Yonchek J, Rodgers KM, Dingman A, et al. Delayed inhibition of tonic inhibition enhances functional recovery following experimental ischemic stroke. J Cereb Blood Flow Metab. 2019;39:1005–1014.

12. Hess G, Aizenman CD, Donoghue JP. Conditions for the induction of long-term potentiation in layer II/III horizontal connections of the rat motor cortex. J Neurophysiol. 1996;75:1765–1778.

13. Chen SX, Kim AN, Peters AJ, Komiyama T. Subtype-specific plasticity of inhibitory circuits in motor cortex during motor learning. Nat Neurosci. 2015;18:1109–1115.

14. Molina-Luna K, Pekanovic A, Röhrich S, Hertler B, Schubring-Giese M, Rioult-Pedotti M-S, Luft AR. Dopamine in Motor Cortex Is Necessary for Skill Learning and Synaptic Plasticity. PLoS One. 2009;4:e7082.

15. Kronenberg G, Balkaya M, Prinz V, Gertz K, Ji S, Kirste I, Heuser I, Kampmann B, Hellmann-Regen J, Gass P, et al. Exofocal dopaminergic degeneration as antidepressant target in mouse model of poststroke depression. Biol Psychiatry. 2012;72:273–281.

16. Alasoadura M, Leclerc J, Hazime M, Leprince J, Vaudry D, Chuquet J. The Excessive Tonic Inhibition of the Peri-infarct Cortex Depresses Low Gamma Rhythm Power During Poststroke Recovery. J. Neurosci. [Internet]. 2024 [cited 2024 Dec 10];44. Available from: https://www.jneurosci.org/content/44/49/e1482232024

17. Saltaji H, Armijo-Olivo S, Cummings GG, Amin M, da Costa BR, Flores-Mir C. Influence of blinding on treatment effect size estimate in randomized controlled trials of oral health interventions. BMC Med Res Methodol. 2018;18:42.

18. Faul F, Erdfelder E, Lang A-G, Buchner A. G*Power 3: a flexible statistical power analysis program for the social, behavioral, and biomedical sciences. Behav Res Methods. 2007;39:175–191.

19. Schallert T, Fleming SM, Leasure JL, Tillerson JL, Bland ST. CNS plasticity and assessment of forelimb sensorimotor outcome in unilateral rat models of stroke, cortical ablation, parkinsonism and spinal cord injury. Neuropharmacology. 2000;39:777–787.

20. Lenth RV. emmeans: Estimated Marginal Means, aka Least-Squares Means [Internet]. 2017 [cited 2024 Aug 22];1.10.4. Available from: https://CRAN.R-project.org/package=emmeans

21. Kuznetsova A, Brockhoff PB, Christensen RHB. lmerTest Package: Tests in Linear Mixed Effects Models. Journal of Statistical Software. 2017;82:1–26.

22. Kassambara A. rstatix: Pipe-Friendly Framework for Basic Statistical Tests [Internet]. 2019 [cited 2024 Aug 22];0.7.2. Available from: https://CRAN.R-project.org/package=rstatix

23. Vickers AJ. The use of percentage change from baseline as an outcome in a controlled trial is statistically inefficient: a simulation study. BMC Medical Research Methodology. 2001;1:6.

24. Alnosaier W, Birkes D. Inner workings of the Kenward–Roger test. Metrika. 2019;82.

25. Joy MT, Carmichael ST. Encouraging an excitable brain state: mechanisms of brain repair in stroke. Nat Rev Neurosci. 2021;22:38–53.

26. Clark TA, Sullender C, Jacob D, Zuo Y, Dunn AK, Jones TA. Rehabilitative Training Interacts with Ischemia-Instigated Spine Dynamics to Promote a Lasting Population of New Synapses in Peri-Infarct Motor Cortex. J. Neurosci. 2019;39:8471–8483.

27. Joy MT, Carmichael ST. Learning and stroke recovery: parallelism of biological substrates. Seminars in neurology. 2021;41:147.

28. Winterbottom L, Nilsen DM. Motor Learning Following Stroke. Physical Medicine and Rehabilitation Clinics of North America. 2024;35:277–291.

29. Murphy TH, Corbett D. Plasticity during stroke recovery: from synapse to behaviour. Nat Rev Neurosci. 2009;10:861–872.

30. Huang L, Lafaille JJ, Yang G. Learning-dependent dendritic spine plasticity is impaired in spontaneous autoimmune encephalomyelitis. Developmental Neurobiology. 2021;81:736–745.

31. Wahl AS, Omlor W, Rubio JC, Chen JL, Zheng H, Schröter A, Gullo M, Weinmann O, Kobayashi K, Helmchen F, et al. Asynchronous therapy restores motor control by rewiring of the rat corticospinal tract after stroke. Science. 2014;344:1250–1255.

32. Allegra Mascaro AL, Conti E, Lai S, Di Giovanna AP, Spalletti C, Alia C, Panarese A, Scaglione A, Sacconi L, Micera S, et al. Combined Rehabilitation Promotes the Recovery of Structural and Functional Features of Healthy Neuronal Networks after Stroke. Cell Reports. 2019;28:3474–3485.e6.

33. Brown CE, Li P, Boyd JD, Delaney KR, Murphy TH. Extensive Turnover of Dendritic Spines and Vascular Remodeling in Cortical Tissues Recovering from Stroke. J. Neurosci. 2007;27:4101–4109.

34. Mostany R, Chowdhury TG, Johnston DG, Portonovo SA, Carmichael ST, Portera-Cailliau C. Local hemodynamics dictate long-term dendritic plasticity in peri-infarct cortex. J Neurosci. 2010;30:14116–14126.

35. Branscheidt M, Hadjiosif AM, Anaya MA, Keller J, Widmer M, Runnalls KD, Luft AR, Bastian AJ, Krakauer JW, Celnik PA. Reinforcement Learning is Impaired in the Sub-acute Post-stroke Period. Neurorehabil Neural Repair. 2025;15459683241304352.

36. Marsh EB, Khan S, Llinas RH, Walker KA, Brandt J. Multidomain cognitive dysfunction after minor stroke suggests generalized disruption of cognitive networks. Brain and Behavior. 2022;12:e2571.

37. Wagner F, Rogenz J, Opitz L, Maas J, Schmidt A, Brodoehl S, Ullsperger M, Klingner CM. Reward network dysfunction is associated with cognitive impairment after stroke. NeuroImage: Clinical. 2023;39:103446.

38. Widmer M, Lutz K, Luft AR. Reduced striatal activation in response to rewarding motor performance feedback after stroke. Neuroimage Clin. 2019;24:102036.

39. Rothman JS, Cathala L, Steuber V, Silver RA. Synaptic depression enables neuronal gain control. Nature. 2009;457:1015–1018.

40. Silver RA. Neuronal arithmetic. Nat Rev Neurosci. 2010;11:474–489.

41. Brown CE, Aminoltejari K, Erb H, Winship IR, Murphy TH. In Vivo Voltage-Sensitive Dye Imaging in Adult Mice Reveals That Somatosensory Maps Lost to Stroke Are Replaced over Weeks by New Structural and Functional Circuits with Prolonged Modes of Activation within Both the Peri-Infarct Zone and Distant Sites. J. Neurosci. 2009;29:1719–1734.

42. Marsh EB, Brodbeck C, Llinas RH, Mallick D, Kulasingham JP, Simon JZ, Llinás RR. Poststroke acute dysexecutive syndrome, a disorder resulting from minor stroke due to disruption of network dynamics. Proceedings of the National Academy of Sciences. 2020;117:33578–33585.

43. Soleimani B, Dallasta I, Das P, Kulasingham JP, Girgenti S, Simon JZ, Babadi B, Marsh EB. Altered directional functional connectivity underlies post-stroke cognitive recovery. Brain Commun. 2023;5:fcad149.

44. Rocha RP, Koçillari L, Suweis S, De Filippo De Grazia M, de Schotten MT, Zorzi M, Corbetta M. Recovery of neural dynamics criticality in personalized whole-brain models of stroke. Nat Commun. 2022;13:3683.

45. Lavin A, Nogueira L, Lapish CC, Wightman RM, Phillips PEM, Seamans JK. Mesocortical dopamine neurons operate in distinct temporal domains using multimodal signaling. J Neurosci. 2005;25:5013–5023.

46. Benoit-Marand M, Ballion B, Borrelli E, Boraud T, Gonon F. Inhibition of dopamine uptake by D2 antagonists: an in vivo study. Journal of Neurochemistry. 2011;116:449– 458.

47. Yang Y, Wang X, Frerking M, Zhou Q. Spine Expansion and Stabilization Associated with Long-Term Potentiation. J. Neurosci. 2008;28:5740–5751.

48. Hayashi-Takagi A, Yagishita S, Nakamura M, Shirai F, Wu YI, Loshbaugh AL, Kuhlman B, Hahn KM, Kasai H. Labelling and optical erasure of synaptic memory traces in the motor cortex. Nature. 2015;525:333–338.

49. Harris KM, Kuwajima M, Flores JC, Zito K. Synapse-specific structural plasticity that protects and refines local circuits during LTP and LTD. Philosophical Transactions of the Royal Society B: Biological Sciences. 2024;379:20230224.

50. Ma Z, Liu H, Komiyama T, Wessel R. Stability of motor cortex network states during learning-associated neural reorganizations. J Neurophysiol. 2020;124:1327–1342.

